# Modeling Mountain Pine Beetle Abundance in Novel Hosts

**DOI:** 10.1101/2024.11.21.624504

**Authors:** Xiaoqi Xie, Micah Brush, Evan Johnson, Catherine Cullingham, Jessica Duffy, Mark A. Lewis

## Abstract

The mountain pine beetle has recently expanded its range into northern and central Alberta, posing an immediate threat to the novel host, jack pine. To date, some experiments suggest that jack pine has limited defensive capabilities despite of the restrictions from the physical environment. In this work, we explore the susceptibility of jack pine compared to the primary host, lodgepole pine, evaluating the risk of potential range expansion into Canada’s boreal forest. We employ a hierarchical model, incorporating environmental and ecological covariates, to examine mountain pine beetle dynamics in a pine forest with lodgepole, hybrid and jack pines. Our results show that pine species significantly influence the probability of being killed, with jack pine being less likely to be infested than lodgepole pine, all else being equal. The hierarchical model demonstrates that beetles perform poorer in jack pine, characterized by a reproduction rate 0.16 times that of non-jack pine. Although jack pine is a suitable host, our results indicate that the number of infestations in jack pine could be lower than in lodgepole pine, with a reduced probability of emerging beetles.

## 1 Introduction

Mountain pine beetle (MPB), *Dendroctonus ponderosae* Hopkins, is a wood-boring insect native in North America, inhabiting most species of pines within its geographical range. In Canada, the range of MPB was historically limited to central and southern British Columbia, and south-western Alberta due to climatic conditions (Taylor & Safranyik, 2003). This species experiences periodic outbreaks, and global warming and the lack of forest management practices resulted in expansion of MPB range in the late 1990s. This outbreak occurred in central British Columbia and then spread into the northern Canadian Rocky Mountains (Aukema et al., 2006; Kurz et al., 2008; Nealis & Peter, 2008; Raffa et al., 2008; Robertson et al., 2009a). Following the expansion, the beetles breached the Rocky Mountain Barrier into Alberta, threatening pine species previously unexposed to MPB (Safranyik & Wilson, 2006; Taylor & Safranyik, 2003).

MPB’s life history helps explain their dynamics. They complete one generation per year, progressing through four life stages: egg, larva, pupa, and adult (Reid, 1962). MPB lays approximately 60-80 eggs per egg gallery in the summer (Safranyik, 1989). The larvae then spend the winter and spring under the bark, emerging as adults the following summer. The adults then search for new hosts, where female beetles release chemicals to attract other beetles to the same tree (Borden et al., 1987). Once beetles successfully infest a host tree, they cut-off the tree’s water supply system, causing it to dry out and turn red the following year (Page et al., 2012). Trees that were killed last year and turn red this year are called ‘red-top’ trees.

The range expansion by MPB increases the vulnerability of pine species not previously within the beetle’s range (Safranyik & Wilson, 2006). In northwestern and central Alberta, lodgepole pine hybridizes with jack pine, *Pinus banksiana* Lamb, a species whose habitat extends from northern Alberta to eastern Canada within Canada’s boreal forests (Rudolph & Laidly, 1990). Jack pine is commonly regarded as closely related to lodgepole pine (Moss, 1949; Rice et al., 2007). In comparison to lodgepole pine, jack pine typically has a shorter lifespan, a shorter height, and thinner phloem (Kenkel et al., 1997). Previous studies have indicated that jack pine could be a suitable host with limited defensive capabilities (Cale et al., 2017; Clark et al., 2014; Cullingham et al., 2011). However, cold winter temperatures can hinder brood development, significantly limiting beetle numbers (Taylor & Safranyik, 2003). In jack pine forest, the cold climate combined with the trees’ thinner phloem, can lead to high winter mortality rates among beetles (Amman, 1976; Chang, 1954; Rosenberger et al., 2017).

The differences between lodgepole and jack pines have raised questions about how the dynamics of MPB will present differently (Bleiker et al., 2023; Colgan & Erbilgin, 2011; Ishangulyyeva et al., 2016). We estimated the statistical effect of pine species on MPB infestations, while considering the impacts from covariates representing geography, temperature, moisture content and nearby beetle pressures. We assessed the relative risk of infesting pines and evaluated the reproduction rate of MPB in jack pine. Overall, this study aims to enhance our understanding of MPB dynamics in jack pine as compared to lodgepole pine.

## 2 Methods

### 2.1 Study area and data preparation

To compare the performance of MPB in lodgepole and jack pines, we selected an area in the central forest of Alberta where we could observe the transition from lodgepole pine to jack pine, going from south to north and west to east as illustrated in Figure 1. We obtained infestation data from aerial surveys, ground surveys, and control programs (Agriculture and Forestry, 2020; Sustainable Resource Development, 2007). Aerial surveys are conducted to thoroughly search for infested trees. Observers in rotary-wing aircraft count red-top trees and nearby current-year killed trees (referred to as green-dead trees) using Global Positioning Systems (GPS) to record their locations. These data are called ‘heli-GPS’ data. Following current year aerial surveys, ground surveys and control programs are done by visiting the identified sites directly. We determined the grid cell size based on the distance between helicopter’s flight lines in aerial surveys. Considering that the distance between two flight lines is approximately 500-1000 meters, we made the grid cell size to be 500 meters. This area contains 229,734 cells, but the exact number of cells detected in each year’s survey varies due to differences in the survey area defined by managers. We obtained data from 2007 to 2020, but we only used data since 2011. As shown in Figure 2, the number of infestations in most years, except for 2009, remained low. The spike in infestations in 2009 is believed to have resulted from a long-distance dispersal from Jasper or British Columbia (Carroll et al., 2017). Therefore, to avoid the influence of this peak year and long-dispersal on our estimations, we used data from 2011 to 2020.

**Figure 1:**
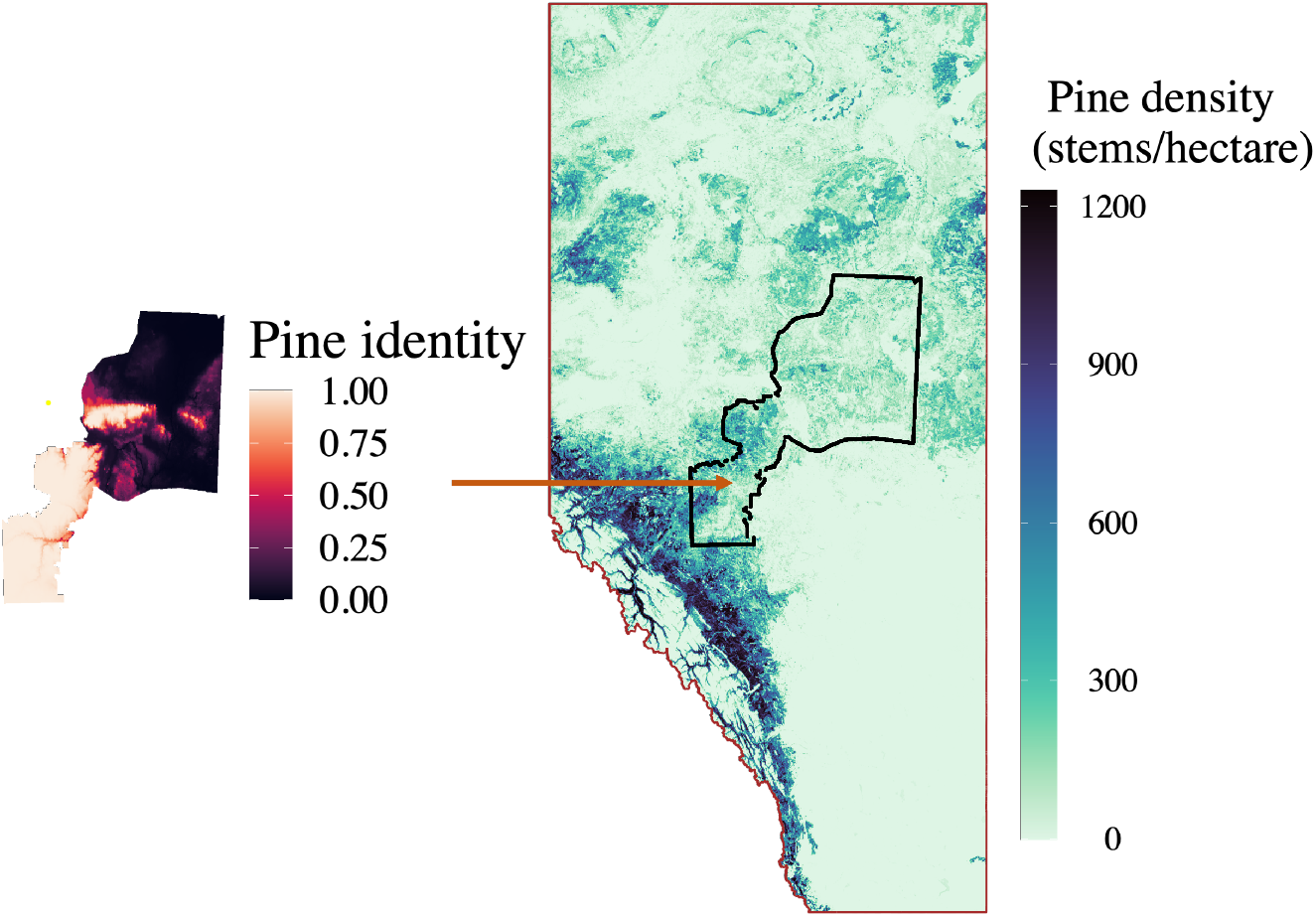
Plots displaying the selected area (highlighted in black) with pine density and identity. The pine identity map has a resolution of 1 km *×* 1 km. The pine identity indicates the species present in each cell. A value less than 0.1 indicates the presence of jack pine. A value greater than 0.9 indicates the presence of lodgepole pine. Values between 0.1 and 0.9 indicate the presence of hybrid pine.

**Figure 2:**
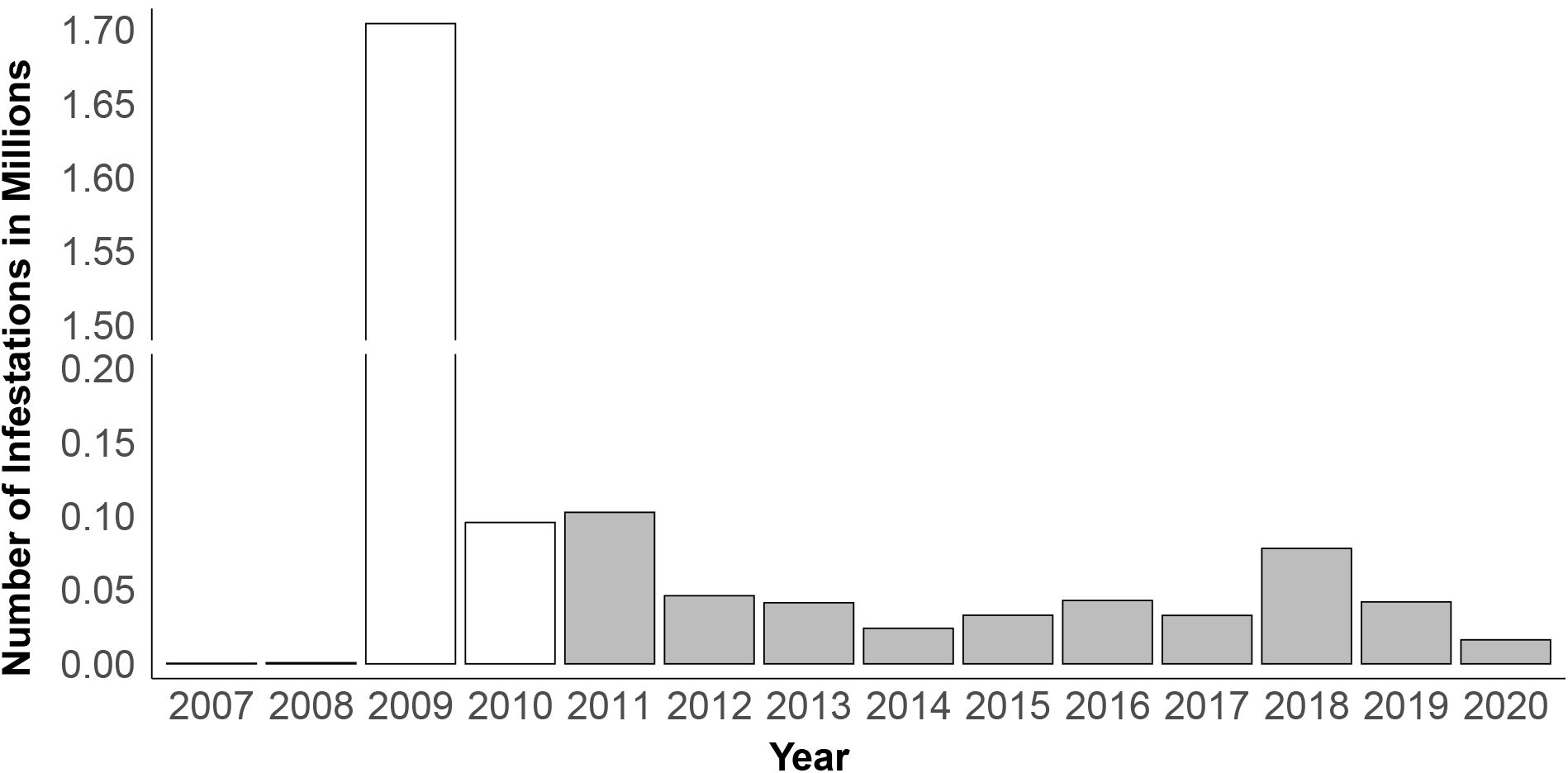
Histogram depicting the observed total infestations within the selected area as shown in Figure 1 over a 14-year period, with counts expressed in millions. Data used in this study are highlighted in grey (2011-2020).

For our model, we considered geographical, climatic, and ecological covariates that have clear biological significance for MPB or their host trees as shown in Table 1. Details regarding covariate selection are provided in Appendix A. We included northness, eastness and slope. Northness and eastness indicate if trees in a cell face north or east, while slope describes the shape of the land. These factors affect sunlight availability, which influences tree height and health. The minimum temperature in summer can restrict beetle dispersal, as beetles typically fly successfully when the temperature is above 19°C and below 41°C (Mccambridge, 1971).

**Table 1:** Overview of model covariates.

| Covariate | Description | Source | Unit |
| --- | --- | --- | --- |
| Northness | Faces north (1) and south (-1). | - | - |
| Eastness | Faces east (1) and west (-1). | - | - |
| Slope | Slope of the cell (Burrough et al., 2015). | - | % |
| Pine identity ($Q$) | Score used to identify pine type. Identifies trees as jack pine if $Q < 0.1$, as lodgepole pine if $> 0.9$, and between is hybrid pine. | (Burns et al., 2019) | - |
| Pine density | Density of jack pine, hybrid pine, and lodgepole pine. The details of the estimation method are provided in Appendix B. | Beaudoin et al. (2014) | stems/cell |
| Age | Average age of the dominant pine trees in each cell. | Beaudoin et al. (2014) | year |
| Minimum temperature in summer | Minimum daily temperature in July and August in the current year. | BioSIM | °C |
| Relative humidity | Average relative humidity from March to May in the current year. | BioSIM | % |
| Wind speed | Average wind speed at 2 meters in July and August in the current year. | BioSIM | m/s |
| Soil moisture index | Average soil moisture index from May to July in the current year. The estimation method is mentioned in Hogg et al. (2013). | BioSIM | % |
| Degree days | Cumulative degree days above 5.5°C from September of the previous year to August of the current year. | BioSIM | °C |
| Overwinter survival probability | Probability of larval survival over the winter during the winter before the current year summer. The estimation method is mentioned in Régnière and Bentz (2007). | BioSIM | % |
| Last year's infestations within a cell | Number of pines killed last year within the cell. | Heli-GPS data | number |
| Dispersal impact within a 4 km radius | Expected number of beetles flying from all directions to the center cell excluding the counted within cell infestations. The details of the estimation method are provided in Appendix B. | Heli-GPS data | number |

The maximum temperature in summer was considered in Appendix A but was excluded here due to its high correlation with degree days. The development of MPB needs at least 833 degree days above 5.6°C a year (Safranyik & Wilson, 2006). Winter survival will impact the beetle population in the following year. We included overwinter survival probability as it calculates the likelihood that a beetle successfully survives until the next spring (Régnière & Bentz, 2007).

Additional covariates were selected for their potential impact on MPB. High wind speeds can help beetles reach remote locations, and last year’s infestations within a cell and dispersal impacts within a 4 km radius estimate beetle pressures from different distances to the center cell. As its name suggests, last year’s infestations within a cell counts the number of infested trees in that cell from the previous year. Dispersal impact within a 4 km radius was estimated from last year’s infestations and represents the expected number of beetles that can come from neighboring areas. Previous studies noted that the influence of past infestations could be significant within at least a 1.25 km radius but less important beyond a 4 km radius (Carroll et al., 2017; Kunegel-Lion & Lewis, 2020). Thus, we estimated the expected number of beetles that emerge from infested trees within a 4 km radius while excluding the center cell. We initially considered that the influence within a cell could vary at distances of 0.75 km, 1.25 km, and 4 km (Kunegel-Lion & Lewis, 2020). Our test results in Appendix A showed that we could integrate the influence within a 4 km radius, excluding the cell, as a single covariate. Details of the derivation of these two covariates are provided in Appendix B.

Besides the above covariates, we considered factors that can affect trees. The availability of pines for beetles is shown by pine density, whereas the age of the pines reflects their vigor and resistance to MPB. Moisture content is considered using relative humidity and the soil moisture index. Relative humidity estimates the moisture content in the air, and the soil moisture index reflects the land’s moisture content (Hogg et al., 2013; Oogathoo et al., 2024).

Pine identity is a special index we included (Burns et al., 2019; Cullingham et al., 2011). Its value, ranging from 0 to 1, represents the pine species. A pine identity value of less than 0.1 indicates that the pine species in this cell is jack pine, while a value greater than 0.9 indicates that the species is lodgepole pine. This score also reflects whether a hybrid pine is closer to lodgepole or jack pine. If the score is between, but closer to 0.1, the pine species in this cell is a hybrid pine that is closer to jack pine. In this study, we used this covariate to measure the comparative performance of beetles in lodgepole and jack pines.

### 2.2 Statistical model

In this study, we investigated the dynamics of MPB in jack pine forests by a hierarchical statistical model, which has been validated and used in another study (Xie et al., 2024). We employed this Zero-Inflated Negative Binomial (ZINB) model to analyze data from 2011 to 2020. This model assumes that the excess zero infestations can result from two reasons: no beetle attacks or failure to kill the host. This model first uses a Bernoulli distribution to determine the probability of presence of beetles, followed by a Negative Binomial model to assess the abundance given that the beetle could present. The probability of zero infestation can come from either the Bernoulli presence model or a zero count in the abundance Negative Binomial model.

We use *Y* as the response variable, representing the number of trees infested in the current year, and let *Z* and *X* be the covariate matrices for the presence and abundance models, respectively. We considered similar covariates in two models, meaning that *Z* equals to *X*. We use *γ* and *β* as the coefficient vectors for the presence and abundance models, respectively. We let *ϕ* as the dispersion parameter in the abundance model. We denote *π* as the probability of zero infestation estimated through the presence model, and *μ* represent the expected number of infestations estimated through the abundance model. The function *f*_Bernoulli_ denotes the probability density function of a Bernoulli distribution. The hierarchical ZINB model is written as:

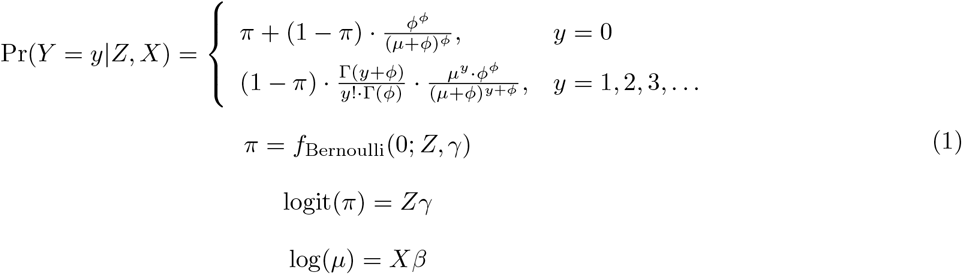

### 2.3 Relative risk of infestation between lodgepole and jack pines

We examined the relative risk as the ratio of the probability of infestations in lodgepole pine to that in jack pine (Nelson et al., 2008). The probability of infestations in either jack or lodgepole pine was estimated by varying the value of the covariate pine identity. We let *y* be the number of infestations in a cell. The probability of infestations in jack pine is estimated as:

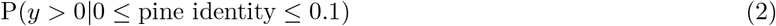

Similarly, we estimate the probability of infestations in lodgepole pine as:

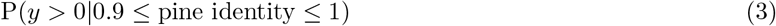

The relative risk is then estimated as:

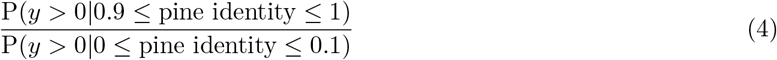

Using the fitted ZINB model, we estimated the probability in different pines by integrating the value of pine identity within the ranges of 0 to 0.1 for jack pine and 0.9 to 1 for lodgepole pine.

### 2.4 Reproduction rate

The reproduction rate in jack pine was estimated as a percentage, indicating how many times less likely or more likely it is compared to the reproduction rate in non-jack pine species, specifically lodgepole and hybrid pines. We did not distinguish between lodgepole and hybrid pines because we want to reduce computational difficulties. We introduced an additional parameter *s* into our ZINB model representing this percentage. We considered the same covariates in the presence and abundance models. This new parameter *s* was inserted into both the presence and abundance models, affecting last year’s jack pine infestations and the dispersal impact from jack pine infestations within a 4 km radius. Each of the two covariates, last year’s infestations within a cell and dispersal impacts within a 4 km radius in one of the two models, was separated into two parts representing its value from jack and non-jack pines.

We use *x* to represent the input covariate vector for a cell and *x*_*l*_ and *x*_*d*_ to represent the covariates of last year’s infestation within a cell and dispersal impact within a 4 km radius, respectively. We use *x*_*l*_*_*_*l*_ and *x*_*l*_*_*_*j*_ to represent last year’s infestations from non-jack pines and jack pines. Since the map of pine identity has a resolution of 500 m, each cell can only contain one pine species. Thus, either *x*_*l*_*_*_*l*_ or *x*_*l*_*_*_*j*_ should be 0. Similarly, we use *x*_*d*_*_*_*l*_ and *x*_*d*_*_*_*j*_ to represent the dispersal impact from non-jack and jack pines. This dispersal impact was estimated by last year’s infestations within a 4 km radius. The values of *x*_*d*_*_*_*l*_ and *x*_*d*_*_*_*j*_ can both be non-zero. We use the same subscripts to highlight the corresponding coefficients. We denote 𝟙 as an indicator function with a bracket indicating the condition for having a value of 1. We have *m* covariates except the intercept. Then, we rewrite *π* and *μ* as:

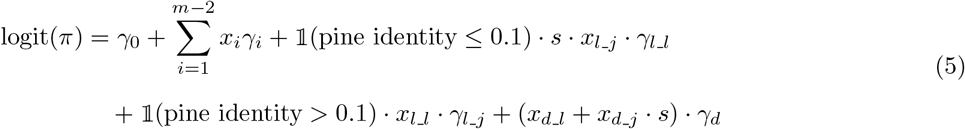

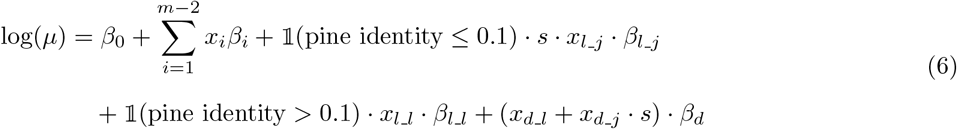

The estimation of the parameter *s* was carried out by minimizing the negative log-likelihood function of ZINB model with the described *μ* and *π* for the presence and abundance models, utilizing the ‘BFGS’ gradient function specified within the optim() function in R.

## 3 Results

Our results demonstrated that lodgepole pine is more susceptible to MPB compared to jack pine. In the fitted hierarchical model, a positive coefficient suggests that an increase in the covariate’s value could raise either the probability or the number of infestations. The probability of infestations is estimated by the presence model, while the number of infestations is calculated by the abundance model. As shown in Figure 3, pine identity was the second largest covariate in the presence model and the largest covariate in the abundance model, marking the potential effect of pine species on beetle’s dynamics. As the value of pine identity increases, the pine species in the cell becomes more similar to lodgepole pine. The large positive coefficients suggest that lodgepole pine has a higher probability of infestation and a greater number of infestations if present. Using Equation (4) and the coefficients in Figure 3, we found that the median value of relative risk is 21.3. By using Equations (5) and (6), the reproduction rate was found to be 0.16. To ensure the reproduction rate remaining unconstrained with values from negative infinity to infinity during the computation, we estimated it in R by its logarithmic value. The log of *s* was -1.833, with a 95% confidence interval of (−1.955, -1.711).

**Figure 3:**
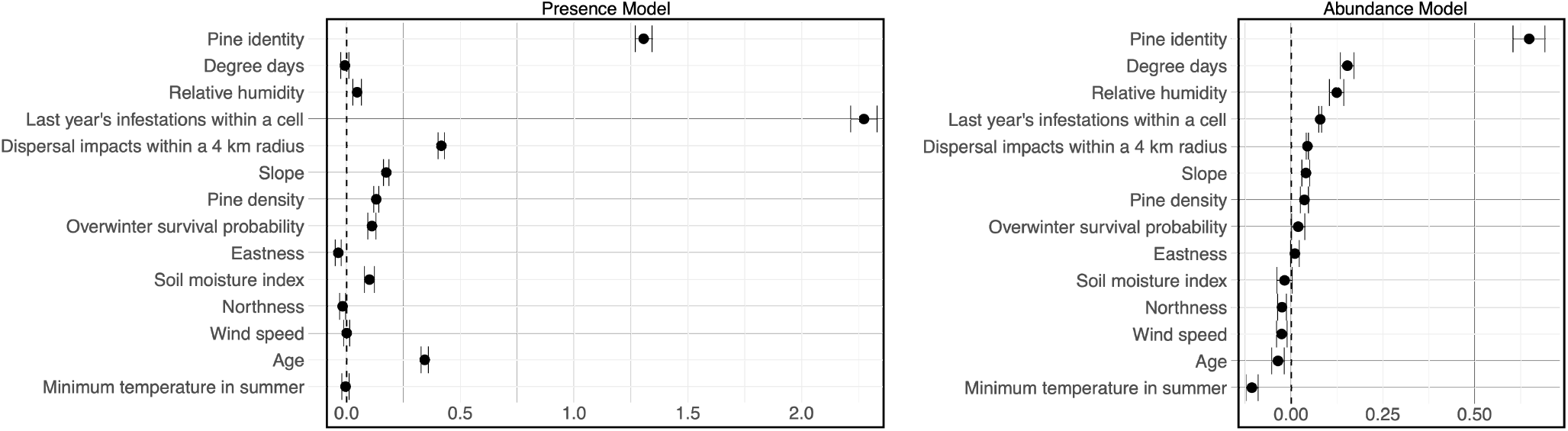
The first plot presents coefficients of standardized covariates from the presence model, and the second plot from the abundance model of the hierarchical model for the specified region (see Figure 1). The coefficient values in two plots are accompanied by their 95% confidence intervals.

As shown in Figure 3, in the presence model, last year’s infestations within a cell and dispersal impacts within a 4 km radius were the largest and forth-largest positive covariates, indicating the significant influence of beetle pressure on the probability of infestations. Results also showed that degree days, wind speed, and minimum temperature in summer were insignificant in the presence model as their confidence intervals included 0. The abundance model demonstrated that last year’s infestations within a cell and dispersal impacts within a 4 km radius played a crucial role for beetles, with higher values suggesting more infested trees. Eastness and soil moisture index were insignificant in the abundance model because their confidence intervals included 0.

## 4 Discussion

The study demonstrated the role of pine species and how environmental and ecological covariates influence MPB population dynamics within a forest containing lodgepole, hybrid, and jack pines. As shown in Figure 3, pine identity is one of the most significant covariates in the presence and abundance models. This suggests that as the pine species in a cell becomes more similar to lodgepole pine, both the probability and number of infestations could increase. The relative risk and reproduction rate presented the same result. The median value of relative risk is 21.3, suggesting that in 50% of the cells, lodgepole pine is at least 21.3 times more likely to be infested compared to jack pine. The reproduction rate is 0.16 when comparing jack pine to non-jack pine. It indicates that the reproduction rate of beetles in non-jack pine is 6.25 times higher than in jack pine when the level of infestations has been consistently low in previous years. Our results, with the consideration of environmental and ecological covariates, suggest that the susceptibility of jack pine to MPB is less than that of lodgepole pine.

Other covariates also showed significance in the model. Last year’s infestations within a cell and dispersal impacts within a 4 km radius were significant in both models, highlighting the influence of beetle pressures on new host trees, agreeing with the previous studies (Carroll et al., 2017; Kunegel-Lion & Lewis, 2020). The positive coefficients for degree days and overwinter survival probability in the abundance model indicated that warmer years and higher survival probability increase the number of infestations. Together, these two covariates suggest that warmer years lead to lower mortality rates benefiting beetles, which is consistent with MPB biology (Régnière & Bentz, 2007; Safranyik & Carroll, 2007).

There are some limitations to this work. By adding data from the peak year 2009, our model places overwhelming emphasis on the significance of dispersal, as indicated by an extremely large positive coefficient.

This large value shows that even a small change in the covariate ‘dispersal impact within a 4 km radius’ results in a significant change in the estimation of infestations. The result suggests that dispersal within a 4 km radius is the most significant covariate, with the impact of changes in other covariates, except last year’s infestations, being negligible, as their coefficient values are much smaller by comparison. To reduce the influence of the peak year data on our model, we used only data from the tail end of the outbreak. Another limitation is the abundance of data. Jack pine thrives in colder locations where beetles struggle to survive the winter, resulting in fewer successful infestations (Aukema et al., 2008; Safranyik, 1978). This cold environment limits the number of infestations in jack pine and reduces the jack pine infestation data for training the model parameter *s*. Additionally, our statistical model assumes that cells are spatially independent, ignoring the spatial autocorrelations that exist among locations. The presence of spatial autocorrelation may lead to an underestimation of the confidence intervals for the model’s parameters (Legendre, 1993). As a result, the significance of covariates with confidence intervals close to zero may vary. However, there is very limited impact on our relative risk value, as both the numerator and denominator would be biased in the same way since they are estimated by the same coefficients.

Our results show a different perspective compared to historical suggestions. Historically, it was believed that jack pine was equally susceptible (Cullingham et al., 2011; Erbilgin et al., 2014), but recent evidence suggests it is less susceptible (Bleiker et al., 2023; Srivastava & Carroll, 2023). We compared the susceptibility of this new host with that of the primary host, presenting a reduced likelihood of infestations, which may differ from conclusions drawn from tree physiology (Cale et al., 2017). When beetles arrive in a forest with both lodgepole and jack pines, they may tend to select lodgepole pines over jack pine. If a jack pine in such a forest is infested, the number of beetles emerging from it could be lower than those emerging from lodgepole pine.

Additionally, our results support Alberta’s current management strategy, which focuses on managing lodgepole pine, as jack pine shows a lower likelihood of infestation. The significance of beetle pressure within a 4 km radius aligns with Alberta’s control program, which emphasizes the necessity of cutting nearby infested trees to reduce infestations in the following year. Our results mention that the influence of one infested tree can be more than 4 km.

Although our results suggest that the risk of further eastward spread is lower than previously anticipated (Burns et al., 2019; Cullingham et al., 2011), climate change introduces uncertainties into our conclusions, as it could significantly raise winter temperatures, increasing beetle survival probability (Hayhoe & Stoner, 2019). Our findings suggest that, compared to lodgepole pine, jack pine has a lower likelihood of infestation when the level of infestations is consistently low, and beetles have a reduced reproduction rate in jack pine. Our conclusions are based on historical environmental and ecological data, which indicated that jack pine forests provide a less favorable environment for beetles. However, ongoing climate change may alter this scenario, placing jack pine at greater risk. Jack pine forests still require attention.

## Supporting information

Appendice

## Acknowledgements

We would like to thank all members of the Lewis Research Group, particularly Kévan Rastello, for his comments on this project. We would also like to thank the members of the TRIA-FoR project for their support. We thank Kelsey Gritter for her help with data wrangling. Special thanks go to the Forestry Division, Government of Alberta, especially David Strauss and Caroline Whitehouse, for their assistance with our data. XX acknowledges the use of ChatGPT for assistance in checking grammar and vocabulary.

## Funding

Funding for this research has been provided through grants to the TRIA-FoR Project to MAL from Genome Canada (Project No. 18202) and the Government of Alberta through Genome Alberta (Grant No. L20TF), with contributions from the University of Alberta and fRI Research (Project No. U22004). MB acknowledges the support of the Natural Sciences and Engineering Research Council of Canada (NSERC), [PDF – 568176 - 2022].

## Competing interests

The authors declare there are no known competing interests.

## Author contributions

Conceptualization: Catherine Cullingham, Evan Johnson, Mark A. Lewis, Micah Brush, Xiaoqi Xie; Methodology: Evan Johnson, Mark A. Lewis, Micah Brush, Xiaoqi Xie;

Formal analysis and investigation: Xiaoqi Xie;

visualization: Xiaoqi Xie;

Software: Xiaoqi xie;

Supervision: Mark A. Lewis;

Writing – original draft: Xiaoqi Xie;

Writing – review & editing: Catherine Cullingham, Evan Johnson, Micah Brush, Mark A. Lewis, Jessica Duffy, Xiaoqi Xie;

## Data availability

Data will be available later.

## Notes

### Competing Interest Statement

The authors have declared no competing interest.

