## Appendice for "Modeling Mountain Pine Beetle Abundance in Novel Hosts"

### List of Figures

1 Plots displaying the selected area (highlighted in black) with pine density and identity. The pine identity indicates the species present in each cell. A value less than 0.1 indicates the presence of jack pine. A value greater than 0.9 indicates the presence of lodgepole pine. Values between 0.1 and 0.9 indicate the presence of hybrid pine. . . . . . 4

2 Histogram depicting the observed total infestations within the selected area as shown in Figure 1 over a 14-year period, with counts expressed in millions. Data used in this study are highlighted in grey (2011-2020). . . . . . 4

3 The first plot presents coefficients of standardized covariates from the presence model, and the second plot from the abundance model of the hierarchical model for the specified region (see Figure 1). The coefficient values in two plots are accompanied by their 95% confidence intervals. 9

### Appendix A Covariate selection

The study of mountain pine beetle (MPB) has been ongoing for a long time, primarily focusing on the development of mechanistic models like the age structure models highlighted in the works of Brush and Lewis (2023), Goodsman et al. (2016), and Heavilin and Powell (2008). These models offer valuable insights into the beetle's long-term dynamics. However, their precision diminishes in short-term analyses, an issue particularly relevant for forest managers who focus on short-term dynamics for planning. To make accurate short-term predictions, advancements in computing power have led researchers to adopt data-driven statistical models. Studies such as those by Kaufeld et al. (2014), Kunegel-Lion and Lewis (2020), Preisler et al. (2012), and Wulder et al. (2006) have implemented regression models to estimate infestations, demonstrating effectiveness in providing short-term predictive visualizations that align more closely with the planning needs of forest managers.

The objective of this appendix was to select covariates that are significant when using considered models. We utilized the area as shown in Figure Appendix A.1 with a cell size of 500 meters. Past work reveals that covariates at different stages can present opposing influences on infestations (Kunegel-Lion & Lewis, 2020). We divided 14 years of data into two stages: outbreak and growth peak declining, as shown in Figure Appendix A.2. We present the significant covariates in the outbreak stage.

Our initial set of covariates was built on Kunegel-Lion and Lewis (2020). Our investigation was mainly guided by the following biological hypotheses:

1. Geographical covariates (northness, eastness, and slope) affect whether beetles are able to infest the stand and the number of trees that succumb to the beetle.

2. Climatic covariates (maximum temperature in summer, minimum temperature in summer, relative humidity, wind speed, soil moisture index, degree days, and overwinter survival probability) affect whether beetles are able to infest the stand and the number of trees that succumb to the beetle.

3. Ecological covariates (pine age, age$^2$, pine height, last year's infestations within a cell, within the adjacent cells, the neighborhood of cells adjacent to those adjacent cells, and weighted influence within a 4 km radius) affect whether beetles are able to infest the stand and the number of trees that succumb to the beetle.

4. There is a quadratic relationship between pine age and resistance (Safranyik et al., 1975).

5. The height derived from an age-height model is significant (see Xie, 2024 for details of the height calculation).

6. The influences of dispersal at local and non-local distances differ.

Elevation was removed because it has a high correlation with degree days. The methods we used to derive covariates are mentioned in Appendix B. Geographic covariates were from the public maps. They were directly interpolated from the public maps. We summarized covariates in Table Appendix A.1.

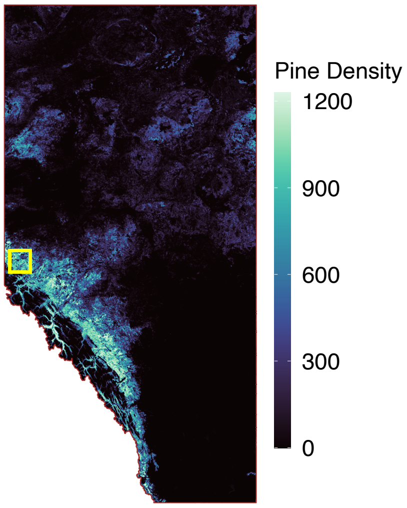


Figure Appendix A.1: Study area selected for covariate selection, highlighted in yellow. Pine density is measured in stems per hectare.

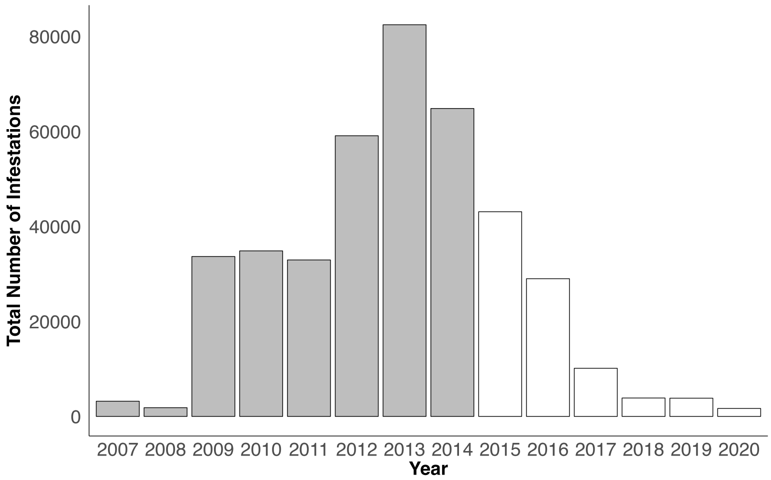


Figure Appendix A.2: Histogram depicting the observed total infestations within the selected area over a 14-year period, split into two stages. The outbreak stage is shown in grey (2007-2014) and the growth peak declining stage is shown in transparent (2015-2020).

| Variables | Description |
|---|---|
| Northness | Face north (1) and south (-1). |
| Eastness | Face east (1) and west (-1). |
| Slope | Slope of the cell (Burrough et al., 2015). |
| Pine density | Pine density (stems/cell; composing of lodgepole pine and jack pine) (Beaudoin et al., 2014). |
| Age | Average age of the leading pines in each cell (Beaudoin et al., 2014). |
| $Age^2$ | Squared age (Beaudoin et al., 2014). |
| Pine height | (Xie, 2024). |
| Maximum temperature in summer | Highest daily temperature in summer in the current year. |
| Minimum temperature in summer | Minimum daily temperature in summer in the current year. |
| Relative humidity | Average relative humidity from March to May in the current year. |
| Wind speed | Wind Speed at 2 meters in July and August in the current year. |
| Soil moisture index | Average soil moisture index from May to July, estimating the water supply for tree growth; current year (%) (Hogg et al., 2013). |
| Degree days | Cumulative degree days above 5.5°C from the September of the previous year to the August in the current year (Safranyik et al., 1975). |
| Overwinter survival probability | Probability of larval survival over the winter; the winter before the current year summer (%) (Régnière & Bentz, 2007). |
| Last year's infestations within a cell | Number of trees infested in the last year within the cell. |
| Last year's infestations within 0.75 km | Number of trees infested in the last year within 0.75 km (as shown in Figure Appendix A.3a). |
| Last year's infestations within 1.25 km | Number of trees infested in the last year within 1.25 km (as shown in Figure Appendix A.3b). |
| Dispersal impacts within a 4 km radius | Expected number of beetles flying from all directions to the center cell excluding the center cell, adjacent cells of the center cells and cells adjacent to the adjacent cells. |
| Infestations (Response variable) | Number of trees infested in the current year. |

Table Appendix A.1: Descriptions of the initial covariates and response variable.

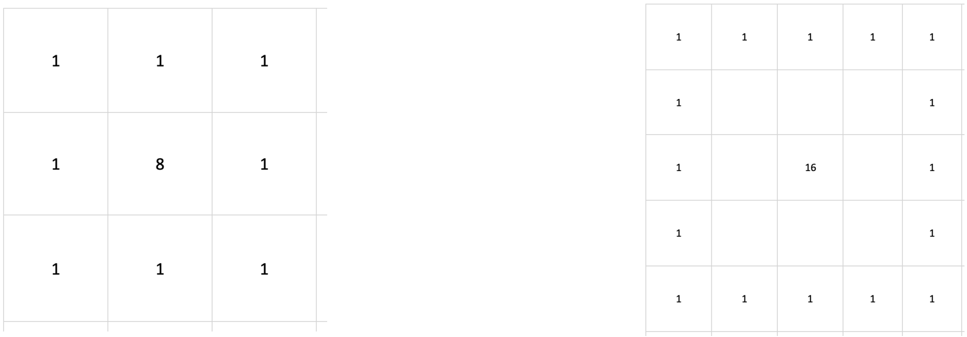


(a) (b)

Figure Appendix A.3: (a) Description of the method we used to count last year's infestations within 0.75 km.(b) Description of the method we use to count last year's infestations within 1.25 km.

We tested the significance of covariates in the model using regularization techniques, which provide a rapid and effective solution for addressing multicollinearity and variable selection (Fan & Li, 2001; Hoerl & Kennard, 1970). We looked for covariates that were significant when considering both issues.

Multicollinearity can significantly impact prediction accuracy, while variable selection helps reduce model complexity. In our dataset, we encountered both data-based multicollinearity, resulting from pure observations, and structural multicollinearity, where covariates use other covariates as input covariates. We wanted to minimize these collinearities, ensuring that covariates provide unique information in the regression model. To achieve this, we made use of Ridge regression (Hoerl & Kennard, 1970). Let $\theta^T$ represent the parameter vector ($\beta$, $\xi$), where $\beta$ contains coefficients of the model, and $\xi$ encompasses all other parameters. Additionally, let $\ell(\theta)$ denote the log-likelihood function. The estimator for Ridge regression, as described in Hoerl and Kennard (1970), is defined as:

$$\hat{\theta} = \text{argmin}_{\theta}(-\ell(\theta) + \lambda_1 \sum_{i=1}^{n} \beta_i^2) \tag{Appendix A.1}$$

where $\lambda_1$ is the tuning parameters of the model and $n$ represents the number of non-intercept coefficients in the model.

Except multicollinearity, we had another concern about model complexity. We wanted to concentrate specifically on the most significant covariates. However, the exhaustive best subset selection method employed in Kunegel-Lion and Lewis (2020) is computationally expensive, as a model with $n$ covariates would result in $2^n$ subsets. To overcome this limitation, we employed a lasso-type penalty for variable selection. A lasso-type penalty prunes small coefficients to zero (Tibshirani, 1996). Initially, we considered three popular lasso-type penalties: Adaptive Lasso, Smoothly Clipped Absolute Deviation (SCAD), and Minimax Concave Penalty (MCP) (Fan & Li, 2001; Zhang, 2010; Zou, 2006). Among these options, we chose SCAD penalty based on the findings of Z. Wang et al. (2016), who demonstrated that SCAD produces less biased estimators while maintaining continuity in model prediction. H. Wang et al. (2007) also noted that theoretically, BIC serves as a better information criterion. Compared to lasso penalty, SCAD penalty assigns greater penalization to small coefficients while still considering the impact of larger coefficients. Let $\theta$ contains all parameters similar to the definition in Ridge regression. The SCAD penalty is defined as follows (Fan & Li, 2001):

$$p_{\lambda_2}(|\theta|) = \begin{cases} \lambda_2|\theta|, & \text{if } |\theta| \leq \lambda_2 \\ \frac{2\delta_1\lambda_2|\theta| - \theta^2 - \lambda_2^2}{2(\delta_1 - 1)}, & \text{if } \lambda_2 < |\theta| < \delta_1\lambda_2 \\ \frac{\lambda_2^2(\delta_1+1)}{2}, & \text{if } |\theta| \geq \delta_1\lambda_2 \end{cases} \qquad \text{(Appendix A.2)}$$

where $\delta_1$ is a free parameter greater than 2 and $\lambda_2$ is the tuning parameter. In our analysis, we used the recommended value of 3.7 for $\delta_1$ (Fan & Li, 2001).

In order to incorporate Ridge and SCAD penalties simultaneously into our model, we assigned distinct weights to each penalty and these weights were set in such a way that their sum equals 1. The objective was to find the optimal balance between two penalties, leveraging the variance-reducing property of Ridge Regression and the variable selection capability of SCAD penalty within a single model framework. Let $\theta^T$ represent the parameter vector ($\gamma$, $\beta$, $\xi$), where $\gamma$ contains coefficients of the presence model, $\beta$ contains coefficients of the abundance model, and $\xi$ encompasses all other parameters. We estimated parameters by minimizing the following equation (Z. Wang et al., 2016):

$$\hat{\theta} = \operatorname{argmin}_\theta - \ell(\theta) + \frac{1}{2}(1-\alpha_1)\lambda_3\sum_{i=1}^{b_1}\gamma_i^2 + \frac{1}{2}(1-\alpha_2)\lambda_4\sum_{j=1}^{b_2}\beta_j^2 + \alpha_1 p_{\lambda_3^*}(\gamma) + \alpha_2 p_{\lambda_4^*}(\beta) \qquad \text{(Appendix A.3)}$$

Here, $\alpha_1$ represents the weight of the SCAD penalty in the presence model, while $\alpha_2$ represents the weight of the SCAD penalty in the abundance model. $\lambda_3$ is the tuning parameter of the Ridge regression in the presence model, whereas $\lambda_4$ is the tuning parameter of the Ridge regression in the abundance model. $\lambda_3^*$ is the tuning parameter of the SCAD penalty in the presence model and $\lambda_4^*$ is the tuning parameter in the abundance model. The variable $b_1$ represents the number of coefficients in the presence model excluding the intercept, and $b_2$ represents the number of coefficients in the abundance model excluding the intercept. We set the free parameters $\delta_1$ in the SCAD penalty as shown in equation 12 to be the suggested number 3.7 (Fan & Li, 2001). To implement the regularization, we utilized the `R package mpath` (Z. Wang et al., 2016). However, the `zipath()` function in this package does not provide automatic support searching for different weights. Therefore, we conducted a grid search to determine the optimal weights. The optimal weights and the tuning parameters were selected based on the smallest BIC value when allowing the function to conduct 100 values for each weight. The tuning parameters were assigned automatically by the `zipath()` function. By using the above methods, the optimal weights of penalties for ZINB model for the outbreak data was calculated to be 1 for both the presence and abundance components. The non-zero coefficients of the standardized covariates corresponding to the hypotheses, along with their 95% confidence intervals were shown in Figures Appendix A.4 and Appendix A.5. We presented covariates including age, age$^2$, height, last year's infestations within a cell, within 0.75 km, 1.25 km and dispersal impacts within a 4 km radius separately from other covariates.

As shown in Figure Appendix A.4, during the outbreak stage, both height and age had coefficients of zero, while age square and dispersal impacts within a 4 km radius had confidence intervals closing to zero in the presence model. The abundance model showed non-zero coefficients for age square, height, dispersal impacts within a 4 km radius, and last year's infestations within 1.25 km, albeit with confidence intervals that included or were close to zero. Regarding the additional covariates in Figure Appendix A.5, northness and eastness lacked significance, and soil moisture index had a coefficient of zero in the presence model during the outbreak stage. In the same stage's abundance model, northness, slope, soil moisture index, degree days, eastness, and pine density were of lesser importance.

To simplify the model, we decided to integrate covariates that include last year's infestations within 0.75 km and 1.25 km, as well as dispersal impacts within a 4 km radius, to be one covariate in our analysis as illustrated in Appendix B.

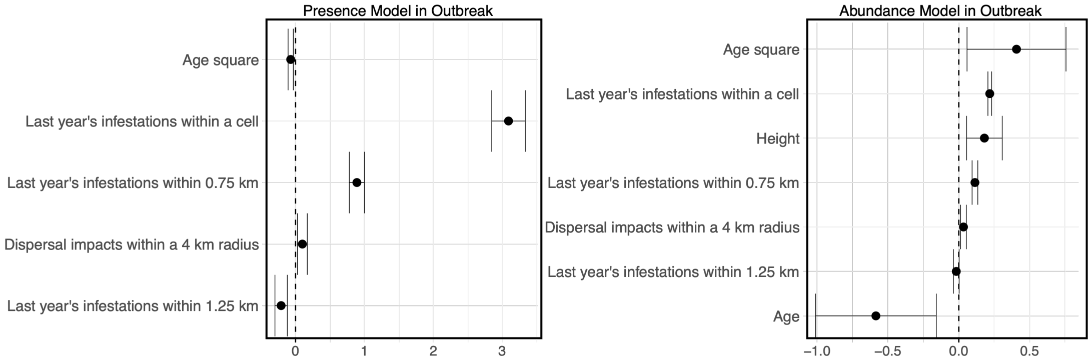


Figure Appendix A.4: Coefficients of age, age$^2$, height, last year infestations within cell, within 0.75 km, 1.25 km and dispersal impacts within 4 km and their 95% confidence intervals in the Presence model in the ZINB model in the outbreak stage (left) and the abundance model in the ZINB model in the outbreak stage (right).

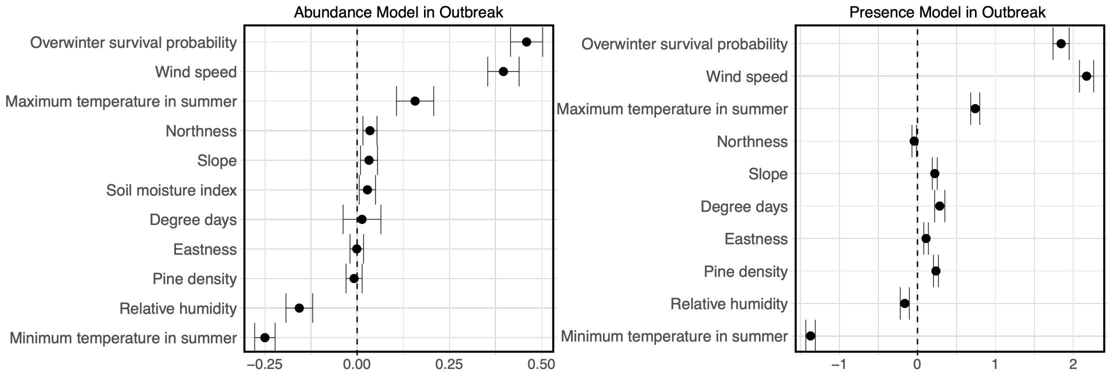


Figure Appendix A.5: Coefficients of covariates not presented in Figure Appendix A.4 and their 95% confidence intervals in the Presence model in the ZINB model in the outbreak stage (left) and the abundance model in the ZINB model in the outbreak stage (right). Covariates not shown in the plots indicate zero values.

### Appendix B Covariate derivation

In this appendix, we show how we derived our covariates.

Our response variable is the number of infested trees killed in the current year. The infestation data were achieved from different surveys. Alberta Environment conducts annual aerial surveys within the designated areas, aimed at thoroughly identifying trees that were infested and died in the previous year. Following the aerial survey, subsequent ground survey and control program are implemented in October and November. Aerial survey and ground survey contain green-dead trees and red-top trees. Green-dead trees are trees killed in the current year and red-top trees are those dead in the previous year. Control program involves cutting and burning the green-dead trees (Government of Alberta, 2023; Sustainable Resource Development, 2007). Thus, the number of infestations in year $t$ is calculated by adding the number of green-dead trees (aerial survey and ground survey) in year $t$ and the number of detected red-top trees (aerial survey) in year $t+1$ to the number of controlled trees (control program) in year $t$.

Our covariates, which included geographical, climatic, and ecological data, were sourced from Alberta Environment, Alberta Vegetation Inventory (AVI), and National Forest Inventory (NFI) (Agriculture and Forestry, 2020; Agriculture, Forestry and Rural Economic Development, 2022; Beaudoin et al., 2014). For climatic covariates not sourced from these three specified places, we used BioSIM to generate the necessary information.

Pine density data was originated from NFI. We did not have explicit data of stems per cell. We had 2011 total volume of all trees measured in volume per hectare, lodgepole pine abundance percentage, and jack pine abundance percentage (Beaudoin et al., 2014). We estimated the pine stems per cell using the developed volume-numbers model from the appendix S1 of Goodsman et al. (2016). The procedure of estimating expected stems per cells is as follows :

1. volume per hectare of lodgepole pine and jack pine = total volume per hectare × (lodgepole pine abundance percentage + jack pine abundance percentage)

2. the expected number per hectare = 17.6 × volume per hectare of lodgepole pine and jack pine× exp(-0.00527 × volume per hectare)

3. the expected number per cell = the expected number per hectare × 25

Since we only had one year volume per hectare value, we then adjusted pine number by subtracting or adding the number of current year infested trees from the recorded year.

We also only had age data for 2011. Thus, we adjusted tree age by subtracting or adding years. We obtained monthly relative humidity, soil moisture index, degree days, wind speed, and yearly overwinter survival probability from BioSIM. We then averaged the relative humidity, soil moisture index, and wind speed over the months of interest and accumulated degree days from September to August of the following year. We directly used the overwinter survival probability from BioSIM, which accounts for the survival probability during the winter preceding the current year's summer.

Last year's infestations within a cell was estimated using the method similar to the response variable except excluding the controlled trees. The observed annual expansion of red-top trees over kilometers led us to hypothesize that the beetles disperse over intermediate distances (Carroll et al., 2017; Robertson et al., 2009b). However, the exact impact of past infestations at different spatial scales on the adjacent cell is not known. Based on Carroll et al. (2017), we proposed that the influence of one infested tree extends up to 4 km. To calculate dispersal impact within a 4km radius, we summed the weighted influence of last year's infested trees within a 4 km radius of the center cell excluding the center cell. Let $D_{t,j}$ represent the number of trees infested in cell $j$ in year $t$ where cell $j$ is within the 4 km radius but is out of the center cell, $\lambda$ represent the reproduction rate of females per stem, and $f_{ij}$ represent the flying influence from cell $j$ to cell $i$, where is from the influence of zone shown in the Chapter 3 of Carroll et al. (2017). The beetle pressure $B_{t,i}$ at time $t$ in cell $i$ was calculated :

$$B_{t,i} = \sum_j D_{t-1,j} * \lambda_j * f_{ij} \qquad \text{(Appendix B.1)}$$

where we fixed the reproduction rate $\lambda$ to 166.7 (females/stem) (Cole, 1969).
